# Trastuzumab deruxtecan kills skin and uveal melanoma in vivo by HER2-independent payload release

**DOI:** 10.64898/2026.10.09.757939

**Authors:** Lisa M Nilsson, Elif Sila Tanaydin, Sofia Stenqvist, David Jansson, Lars Ny, Jonas A Nilsson

**Affiliations:** Sahlgrenska Center for Cancer Research, Departments of Surgery, Institute of Clinical Sciences, Sahlgrenska Academy at University of Gothenburg and Sahlgrenska University Hospital; Sahlgrenska Center for Cancer Research, Departments of Oncology, Institute of Clinical Sciences, Sahlgrenska Academy at University of Gothenburg and Sahlgrenska University Hospital

**Keywords:** antibody-drug conjugate, trastuzumab deruxtecan, melanoma, uveal melanoma, HER2, cathepsin L

## Abstract

**Background:** HER2-directed CAR-T cells can eradicate melanoma despite low HER2 expression. We therefore asked whether melanoma would also respond to trastuzumab deruxtecan (T-DXd), an antibody-drug conjugate targeting HER2.

**Patients and methods:** T-DXd was evaluated in several patient-derived xenograft (PDX) models and as a compassionate use in one patient with metastatic uveal melanoma. We compared T-DXd with another ADC, trastuzumab emtansine (T-DM1), and examined ADC sensitivity and trastuzumab uptake in vitro, and expression of HER2 and cathepsin L (CTSL) in xenografts.

**Results:** T-DXd inhibited all subcutaneous PDXs, while liver models were less sensitive. T-DM1 had limited activity. Unexpectedly, parental and *ERBB2*-knockout melanoma xenografts showed similar initial responses to T-DXd, although knockout tumors regrew earlier. Tumor-derived CTSL was detected outside melanoma cells in the xenografts. Melanoma cells were largely resistant to both ADCs in vitro and showed little trastuzumab uptake compared to HER2-amplified breast cancer cells.

**Conclusions:** T-DXd has a substantial HER2-independent component of activity in the tested melanoma xenografts. Presence of extracellular CTSL, an enzyme known to cleave T-Dxd, provide a possibility that linker cleavage and payload release can occur outside of the tumor cells.

**Highlights:**

– T-DXd caused marked regressions in four subcutaneous uveal melanoma PDXs, prompting treatment of one matched patient.
– The matched patient and orthotopic liver models were less responsive, revealing strong site-dependent activity.
– ERBB2 knockout does not ameliorate therapeutic effects in vivo but correlate with presence of extracellular tumor-derived CTSL.
– Cultured melanoma cells did not exhibit trastuzumab uptake and were not sensitive to T-Dxd or T-DM1.

## Introduction

Human epidermal growth factor receptor 2 (HER2), also known as ErbB2, is a member of the EGFR family of transmembrane tyrosine kinase receptors and plays a pivotal role in regulating cell proliferation, survival, and differentiation. Overexpression or amplification of *ERBB2* encoding HER2 has been identified in several malignancies, including breast, gastric, bladder, and lung cancers, where it is often associated with aggressive tumor behavior and poor prognosis (1). HER2 alterations, including mutations and amplifications, activate downstream signaling pathways such as MAPK/ERK and PI3K/AKT/mTOR, promoting tumor growth and survival (2).

Therapeutic advances have led to the development and approval of multiple HER2-targeted agents, including monoclonal antibodies (e.g. trastuzumab), tyrosine kinase inhibitors, and antibody-drug conjugates (ADCs) such as trastuzumab emtansine (T-DM1) and trastuzumab deruxtecan (T-Dxd). ADCs have particularly transformed the treatment landscape by combining targeted antibody specificity with potent cytotoxic payloads, enabling selective tumor cell killing while limiting systemic toxicity (3). They have also shown efficacy beyond breast and gastric cancers, with HER2 expression detected in a subset of other epithelial malignancies, suggesting broader therapeutic potential (4).

HER2-directed chimeric antigen receptor T cells (CAR-T cells) have shown activity in solid tumors, including sarcoma (5,6). We previously found that the comparatively low HER2 expression in cutaneous and uveal melanoma was sufficient for HER2 CAR-T-cell activity (7) and that HER2-specific CARs introduced into tumor-infiltrating lymphocytes retained therapeutic activity in mouse xenograft models (8). We therefore hypothesized that melanoma could be susceptible to HER2-directed ADCs.

The aim of this study was to determine if uveal and skin melanoma were sensitive to HER2 targeting ADCs. We show that while melanoma xenografts are sensitive to trastuzumab deruxtecan (Enhertu, T-Dxd), this appears largely to be independent of HER2 expression.

## Patient and methods

All experiments were approved by the human research ethics committee (approval numbers 2018-44 and 2020-6916) and the animal ethics committee in Gothenburg (approval numbers 4684-23 and 7146/26). The 1-902 patient-derived xenograft (PDX) mouse model was developed as previously described (9) from a biopsy of a patient with liver metastases of uveal melanoma, that had exhausted all available treatments and trials (Figure 1A). T-Dxd (Enhertu) and T-DM1 (Kadcyla) used in the patient and in mice were from the hospital pharmacy. 6-10 weeks female mice were used in this study. Experimental details, including group sizes, randomization, dosing, routes, endpoints, assay conditions and knockout procedures, are provided in the legends of the figures or supplementary figures.

**Figure 1.**
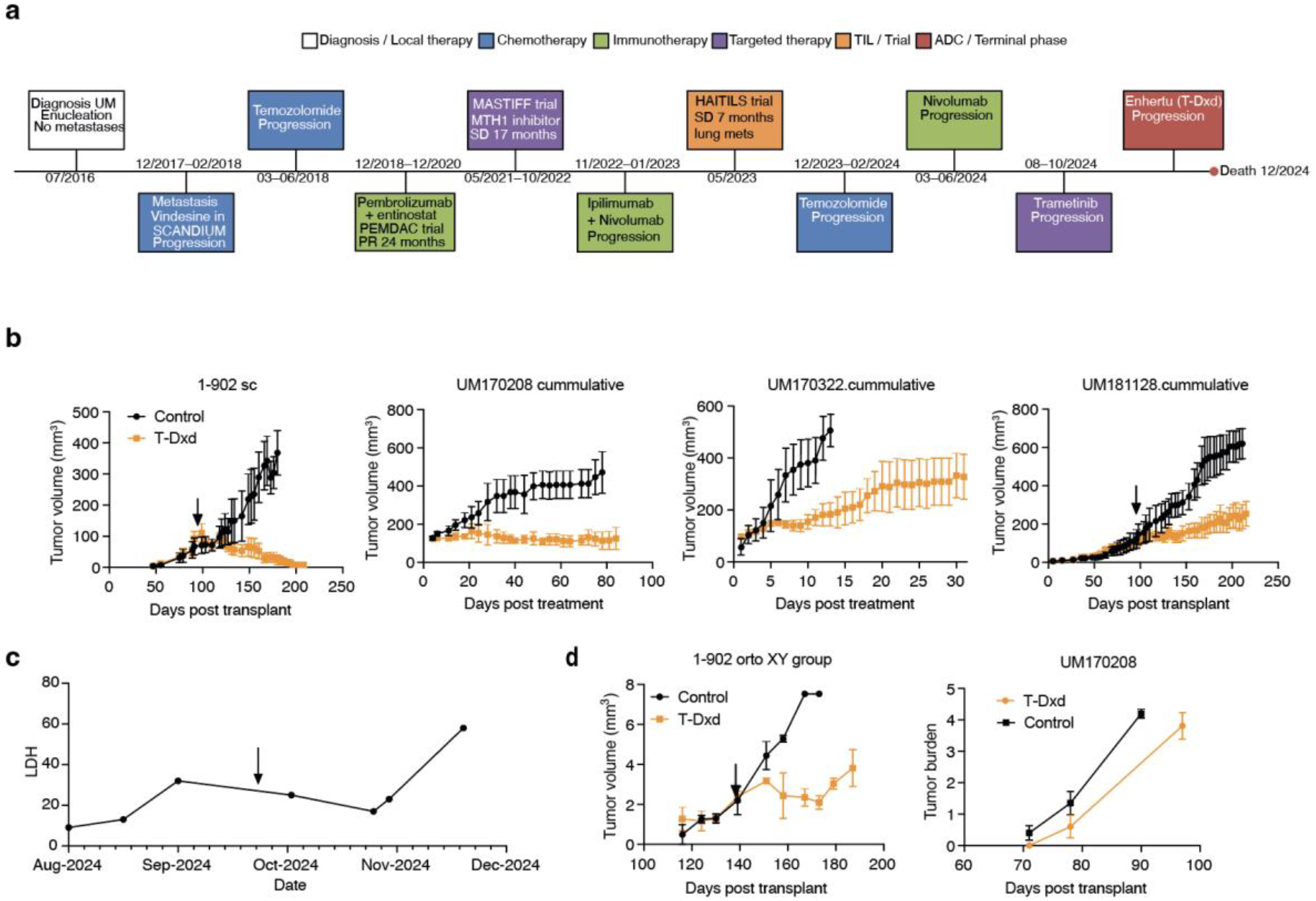
Uveal melanoma PDX models respond to T-Dxd. **a**) Schematic description of the disease progression and treatments of patient 1-902 who participated in four clinical trials at Sahlgrenska University Hospital. In the HAITILS trial, a small part of the tumor resection used for generation of tumor-infiltrating lymphocyte (TIL) therapy was transplanted into the flank of a NOD/SCID/IL2 receptor gamma (NOG) mouse (Taconic) to generate a PDX model. **b**) Mice carrying serially transplanted PDX tumors originating from biopsies of patient 1-902 and three other patients with metastatic uveal melanoma were randomized to either receiving PBS (n=4-5) or T-Dxd (5 mg/kg, n=4-5) once weekly. Subcutaneous tumor growth was monitored by caliper measurements. **c**) After observing tumor growth inhibition by T-Dxd in the 1-902 PDX model (**b**), the corresponding patient was treated with two cycles of T-Dxd (Enhertu, 100 mg). LDH was monitored and upon elevation, treatment was stopped and patient received palliative care. **d**) Orthotopic PDX models carrying liver metastases of indicated PDX tumors were treated with 5 mg/kg T-Dxd when tumors were measurable by ultrasound (Figure S1). Tumor burden was calculated based on the sum of target lesions.

## Results

Mice carrying either the 1-902 PDX tumor from a patient in the HAITILS trial (NCT04812470, Figure 1A), or the UM170208, UM181128 and UM170322 PDX tumors (10), were treated with T-Dxd. All mice responded, with 1-902 and UM170208 exhibiting a complete response (Figure 1B). Because of the response, the 1-902 patient was offered compassionate use of T-Dxd (Enhertu) after progressing in the HAITILS trial (11) and failing subsequent treatments. Only a moderate effect of the drug on tumor marker LDH was seen in the patient (Figure 1C), raising the possibility the high tumor burden or lack of activity of T-Dxd in liver countered the clinical benefit. To address the latter, we generated liver metastasis models from the 1-902 and UM170208 PDX models and treated them with T-Dxd. The 1-902 liver metastasis PDX model had a brief reduction in tumor burden, as assessed by ultrasound, but UM170208 did not (Figure 1D and Figure S1A). Quantification of Ki67 clearly demonstrated a reduction in proliferation in the UM170208 subcutaneous PDX model but not in liver (Figure S1B).

To address more broadly HER2 as a target for ADC in melanoma we treated PDX models with trastuzumab emtansine (T-DM1, Kadcyla). Surprisingly, there was a limited efficacy of T-DM1 in the UM170208 PDX model (Figure 2A) and in combination with T-Dxd the therapeutic effect was worse than with T-Dxd alone. Since expression of HER2 was very low in the melanoma PDX models (Figure 2B) we reasoned HER2 expression might be limiting to allow binding of two targeted agents. Treatment of a HER2 positive pancreatic adenocarcinoma PDX model from patient Pac06 (12) demonstrated antagonism between T-Dxd and T-DM1, which associated with a different HER2 expression pattern between untreated (membranous) and ADC-treated (scattered) tumor cells (Figure S2A-B).

**Figure 2.**
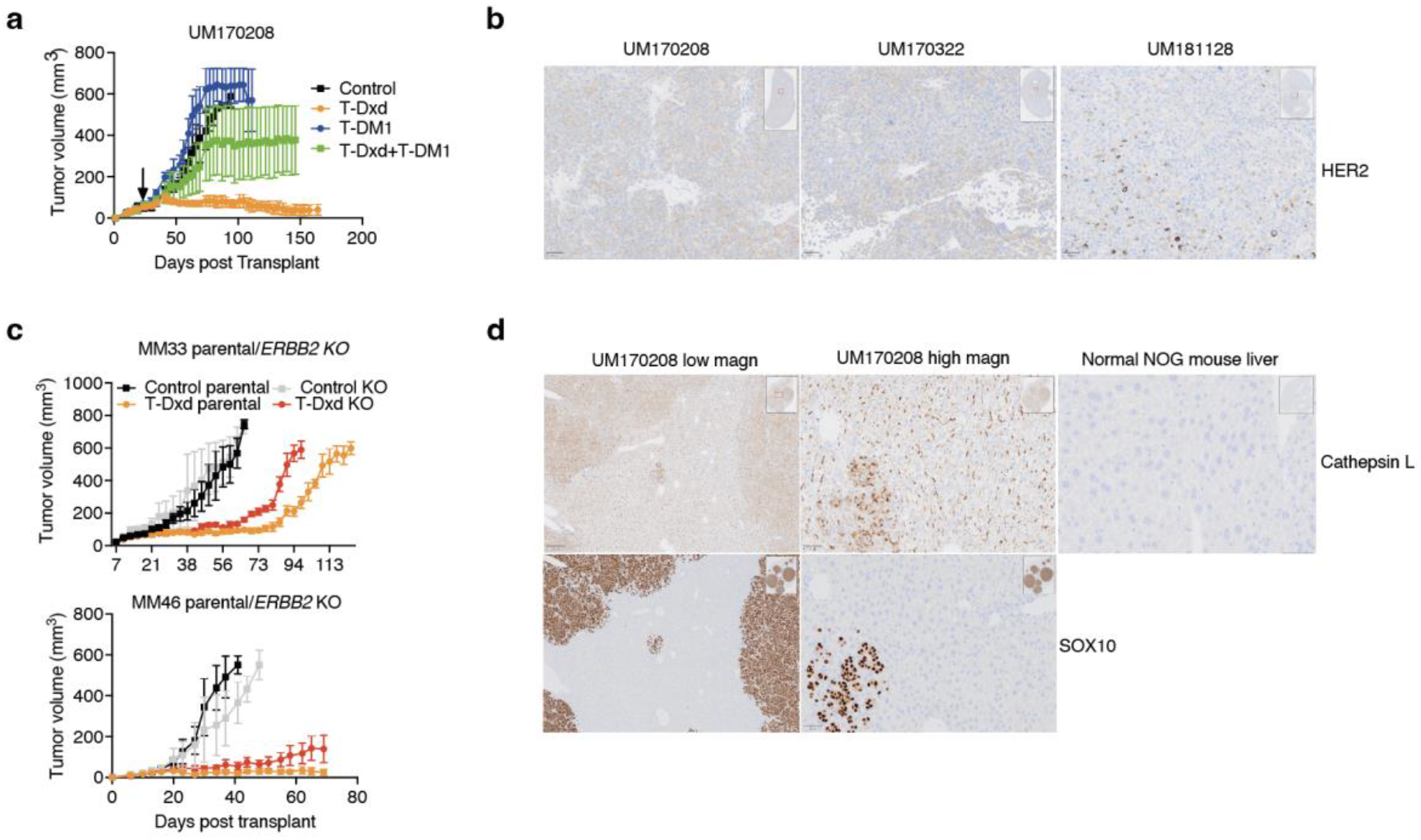
Therapeutic efficacy of T-Dxd correlates with cathepsin L expression but not HER2 expression. **a**) UM170208 PDX model was treated with PBS or 5 mg/kg T-Dxd (intravenously), T-DM1 (Kadcyla, intraperitoneally) or a combination, once weekly. Tumor growth was monitored by caliper measurement. **b**) Immunohistochemistry (IHC) of formalin fixed paraffin embedded tumors of indicated uveal melanoma PDX tumors. Tumors were sectioned (4 µm) and sections were transferred to glass slides and stained using a human HER2 antibody (Dako) in an autostainer. **c**) Patient-derived cutaneous melanoma cells MM33 and MM46 were subjected to CRISPR/Cas9-mediated knockout of *ERBB2* (encoding HER2, Figure S3). Parental and *ERBB2* knockout cells were transplanted into ten NOG mice. When tumors reached ∼50 mm3, mice were randomized to receive PBS or T-Dxd (5 mg/kg). Tumor growth was monitored by caliper measurement. **d**) IHC of UM170208 PDX liver metastases and a normal mouse liver stained with cathepsin L or melanoma marker SOX10 (both Cell signaling) antibodies.

We have previously shown that both uveal and skin melanoma are sensitive to HER2 CAR-T cells (7). We thus xenotransplanted two metastatic skin melanoma cell lines, MM33 and MM46, which previously were shown to be sensitive to HER2 CAR-T cells (7), but which can be made resistant to HER2 CAR-T cells by CRISPR:Cas9 mediated knockout of *ERBB2* (Figure S3A). Both MM33 and MM46 skin melanoma xenografts responded to T-Dxd but unexpectedly, initial response *in vivo* was identical in mice carrying either the parental or HER2 knockout cells (Figure 2C). Resistance development was earlier in mice carrying HER2 deficient melanoma cells, but parental cells also relapsed (Figure 2C). Relapsed cells were resistant to T-Dxd when injected into new mice (Figure S3B).

To investigate if expression of enzymes like cathepsin L (CTSL) could be involved in the HER2 independent therapeutic effects of T-Dxd in melanoma PDX models, we performed immunohistochemistry on mouse PDX models. Since the antibody used was human specific, we could exclusively detect tumor-made CTSL. Intriguingly, the enzyme was not only present inside melanoma cells but also outside of the SOX10 positive melanoma cells (Figure 2D, subQ IHC vs orto). This could explain why T-Dxd could partly be working in a HER2-indpendent manner.

Cathepsin L is known to require an acidic environment, such as the lysosome or extracellularly in hypoxic or metabolically active tumors. To assess the ADCs *in vitro* where pH is neutral, we compared parental and *ERBB2* knockout melanoma cells with the *ERBB2* amplified breast cancer cell line AU565. As expected, AU565 cells were very sensitive to T-DM1 and T-Dxd at 5 days with IC50 of less than 0.05 µg/ml (Figure 3A). On the other hand, the MM33 cutaneous and 92-1 uveal melanoma cells were markedly resistant. The lack of effect in vitro in melanoma correlated with a lack of antibody internalization, as assessed by uptake of an endosome pH-sensitive dye labelled trastuzumab (Figure 3B and 3C). Collectively these findings suggest that T-Dxd does not exhibit therapeutic effects in melanoma in vivo via HER2. The drug is not taken up and does not kill melanoma cells in vitro, but it kills both HER2-proficient and HER2-deficient melanoma in vivo.

**Figure 3.**
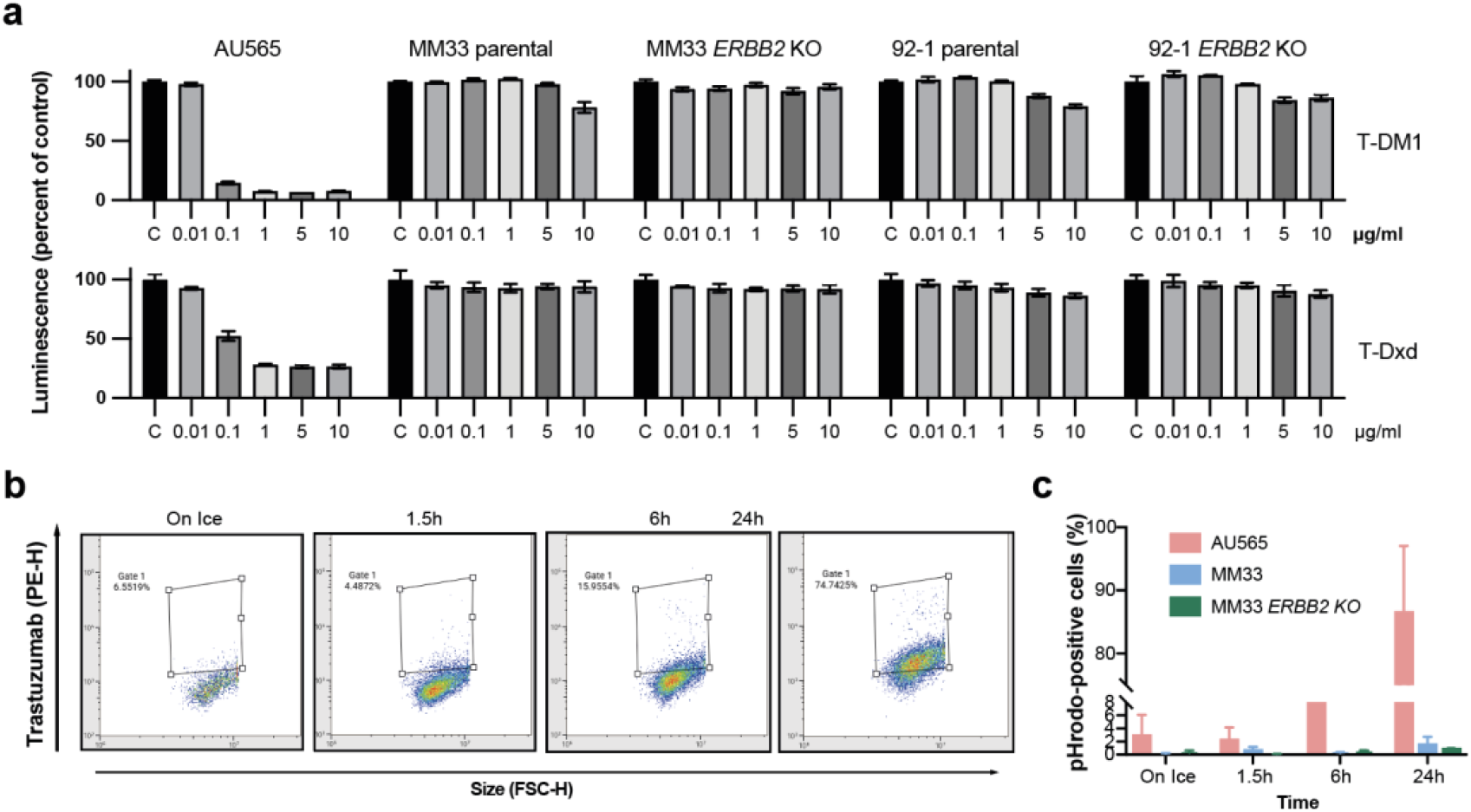
Melanoma cells are resistant to HER2 ADC and are defective in uptake of Trastuzumab-conjugates. **a**) AU565 breast cancer cells, parental or *ERBB2* knockout MM33 cutaneous melanoma cells and parental or *ERBB2* knockout 92-1 uveal melanoma cells were cultured in the presence of increasing concentration of T-DM1 or T-Dxd for 5 days. Cell-titer-glo (Promega) was added and viability was measured using a luminometer. **b-c**) AU565 and MM33 cells were treated with 1 µg/ml Trastuzumab labelled with the pH sensitive dye pHrodo-Red (Invitrogen) on ice. Cells were shifted into 37°C for indicated time points and were then analyzed by flow cytometry. Shown in **b** is the gating and in **c** the average uptake from three independent experiments.

## Discussion

The broad success of T-Dxd (Enhertu) in the clinic, including superior activity to T-DM1 (Kadcyla) in the randomized controlled phase 3 DESTINY trial (13), has sparked an active investigation into just how low HER2 expression is sufficient to still see potent anti-tumoral responses. Although concepts of low and even ultra-low HER2 expression and bystander effects have provided a mechanism (3), alternative explanations could be at play. Recently, a preclinical study demonstrated that cathepsin L (CTSL), an enzyme normally confined to the lysosome, could mediate payload release in the extra-cellular space of tumors (14). Although the authors did not knock out HER2, but instead used cell lines expressing high or undetectable HER2, the data are very much aligned with ours. For instance, both studies demonstrate that HER2 low expressing cells are insensitive to in vitro culture with T-DM1 and T-Dxd.

To investigate if extracellular payload relase via CTSL could be involved in the anti-tumoral effect of T-Dxd in melanoma PDX models, we performed immunohistochemistry on mouse PDX models. Since the antibody used was human specific, we could readily observe tumor-made CTSL in the TME and were intrigued to find the enzyme present outside of the SOX10 positive melanoma cells. While this does not provide casuality, our data along with the deeply mechanistic paper of Tsoa et al. (14), suggest that HER2 expression may be a less informative biomarker of response to T-Dxd than CTSL. Indeed, CTSL expression can vary tremendously in tumors (15–17) and the CTSL gene is also under control of the hypoxia inducible factor Hif1a (18).

Extracellular release of topoisomerase inhibitors by T-Dxd spreading systemically may contribute to side effects like interstitial lung disease, pneumonitis, and acute respiratory distress syndrome. Notably, this side-effect is shared by other Dxd containing ADCs (19). The silver lining is however that CTSL is higher in tumors than in normal tissue. More research into optimizing linkers to ensure increased tumor-selectivity may improve the therapeutic window for T-Dxd and other Dxd containing ADCs. But at this point, focusing on improving methods to detect ultralow expression of HER2 (20) may prove to be futile and unnecessary.

## Acknowledgements

We thank Carina Karlsson for providing technical support. Grant support came from Cancerfonden, Familjen Erling Persson, Vetenskapsrådet, Sjöbergstiftelsen, Västra Götaland Regionen ALF grant to J.A.N, and from Assar Gabrielsson fond to E.S.T.

## Contributions

J.A.N conceptualized and designed the study with L.M.N. providing input and discussions. L.M.N, E.S.T. and S.S performed experiment, analyzed and supported data. D.J. and L.N. conducted patient follow up. J.A.N. interpreted the data, drafted illustrations and wrote the manuscript with input from all the authors. All authors discussed the results.

## Supplementary Figures

**Figure S1.**
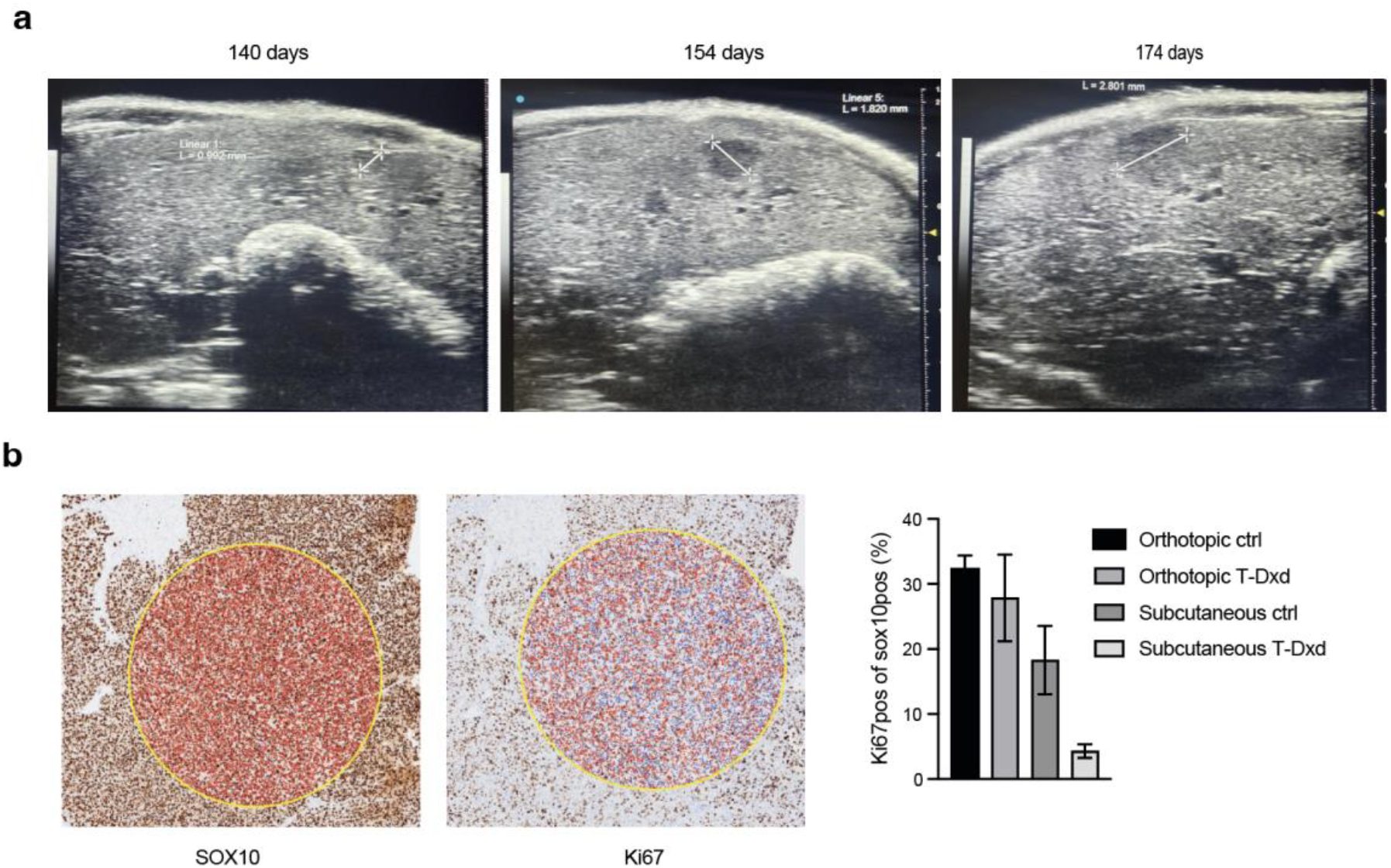
Liver metastases are less responsive to T-Dxd than subcutaneous tumors. **a**) Images of liver metastases in a Vevo 3100 ultrasound instrument (FUJIFILM VisualSonics) showing progression of a target lesion. **b**) Immunohistochemistry using Ki67 antibody (Dako) and melanoma specific marker SOX10 (Cell signaling). *Left* Representative field of views used in the quantification using the imaging software QuPath. *Right* Data obtained from five fields of views of three tumors per treatment group.

**Figure S2.**
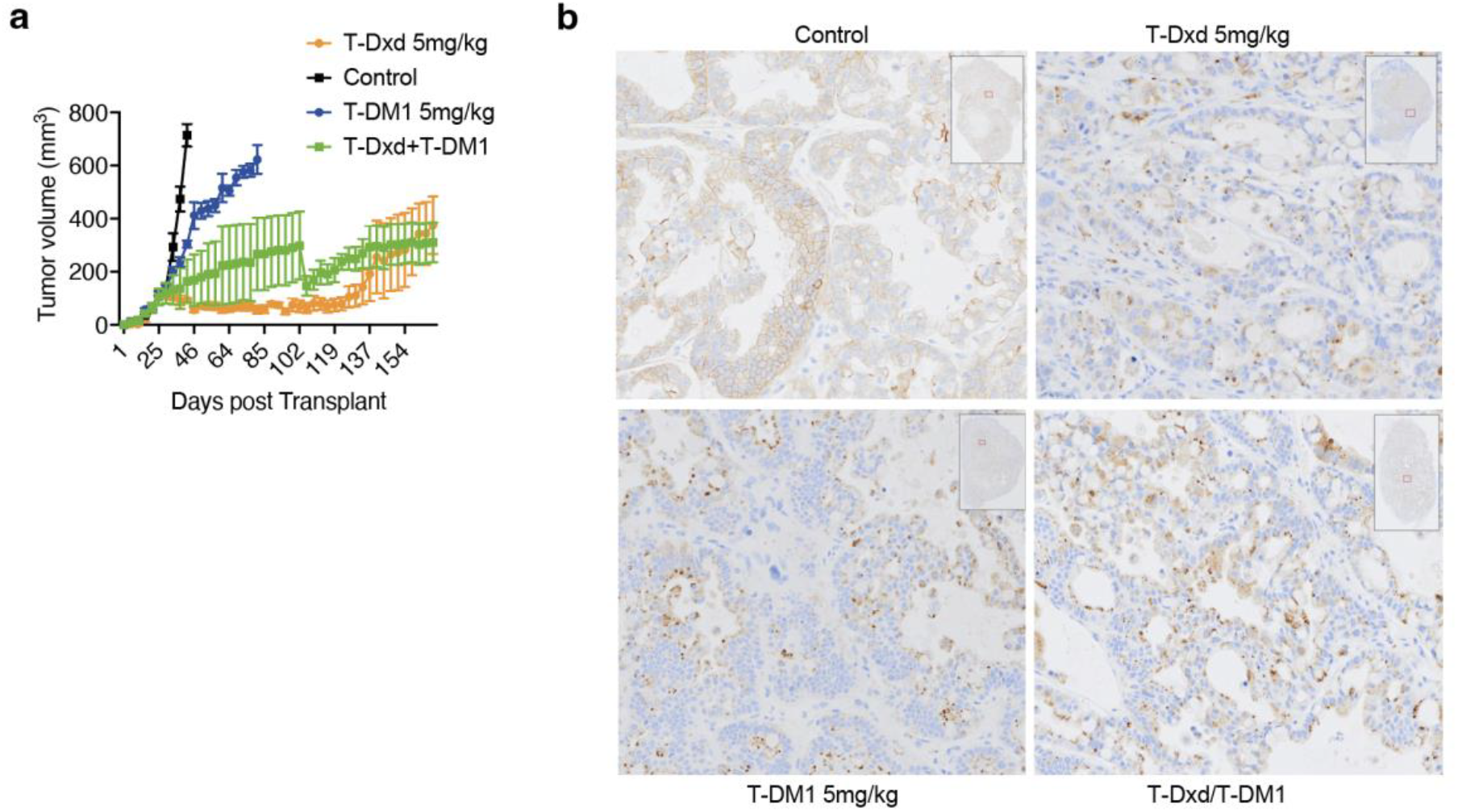
Effect of T-Dxd and T-DM1 in the Pac06 pancreatic cancer PDX. **a**) Tumor bearing NOG mice was treated with PBS or 5 mg/kg T-Dxd (intravenously), T-DM1 (intraperitoneally) or a combination, once weekly. Tumor growth was monitored by caliper measurement. The drop in tumor volume around 100 day was caused by an omission of treatment with Kadcyla. **b**) Immunohistochemistry (IHC) of formalin fixed paraffin embedded tumors from mice treated with indicated drugs using an HER2 antibody.

**Figure S3.**
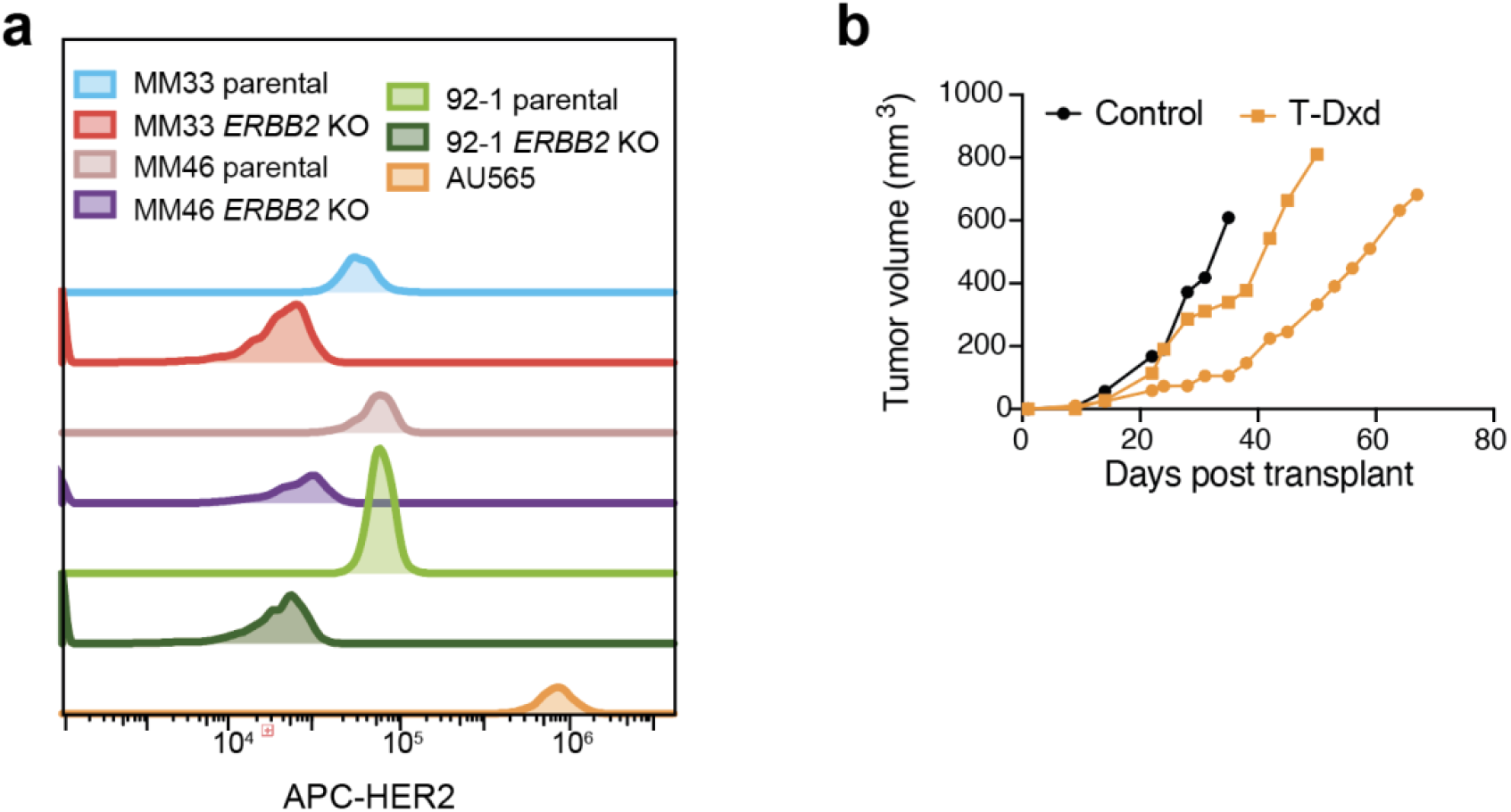
CRISPR:Cas9 mediated knockout melanoma cells lack expression of HER2. **a**) MM33, MM46, 92-1 and AU565 cells cells were grown in RPMI containing 10% fetal bovine serum, penicillin, streptomycin and glutamax (Invitrogen). A guide RNA targeting *ERBB2* (7) was complexed with tracRNA and Cas9 (IDT technologies) and transfected into the MM33, MM46 and 92-1 cells using CRISPRMAX (Invitrogen). Cells were negatively sorted in a cell sorter and expanded. Shown here is a flow cytometry analysis of expanded cells, where expression of HER2 was compared to HER2 amplified breast cancer cell line AU565. **b**) Tumors of xenografted *ERBB2* knockout MM33 relapsing after a complete response after T-Dxd treatment (Figure 2c) were xenografted into new NOG mice. Tumor response to T-Dxd (5 mg/kg) was monitored using caliper measurement.

